# Spatio-temporal distribution patterns and ecological correlates of new mammal records in China

**DOI:** 10.1101/2025.02.09.637225

**Authors:** Chenchen Ding, Jiale Ding, Huijie Qiao, Zhigang Jiang, Zhiheng Wang

**Affiliations:** Institute of Ecology and Key Laboratory for Earth Surface Processes of the Ministry of Education, College of Urban and Environmental Sciences, Peking University; College of Geographical Sciences and Tourism, Xinjiang Normal University; Institute of Zoology, Chinese Academy of Sciences

**Keywords:** biodiversity inventory, mammals, new occurrence records, species distribution, Wallacean shortfall, conservation planning

## Abstract

Species’ geographic distributions are central to research in biogeography, macroecology, and conservation biology. However, incomplete or inaccurate knowledge about their spatiotemporal ranges—known as the Wallacean shortfall—hampers our understanding of biodiversity patterns and processes. In this study, we systematically reviewed 192 papers that reported new distribution records of China’s mammals from 2001 to 2023, covering 150 species in 26 families across 7 orders. We examined the taxonomic, spatiotemporal, and biogeographic characteristics of these newly recorded species. We used Bayesian Phylogenetic Generalized Linear Mixed Models (BPGLMM) to assess how intrinsic traits and extrinsic factors influenced the likelihood of discoveries and partial Poisson regressions to evaluate how species-level attributes, environmental and socio-economic factors shaped the number of new occurrence records across provinces. Our results showed that Chiroptera (n=69), Eulipotyphla (n=26) and Rodentia (n=23) had the highest number of new records. Among provinces, Yunnan (n=31), Guangdong (n=22), and Xizang (n=18) yielded the most newly recorded species. The amount of annual discoveries varied and peaked between 2017 and 2021. The Western Mountain and Plateau (n=39) and the East Hilly Plain (n=33) zoogeographic subregions had the greatest number of newly recorded species, including two new records extending into historically unrecognized zoogeographic realms. Notably, 61 (40.7%) species were found to extend towards the north or northeast of their known ranges, and 32 (21.3%) eastward, mainly due to the sampling bias. Smaller-bodied species and those with broader habitat ranges are more likely to yield new records, and the total number of new records was positively correlated with regional species richness and current survey efforts (r > 0.5, p < 0.05). These findings expand the known distributions of China’s mammals and provide essential data for mitigating the Wallacean shortfall. They underscore the urgent need for intensified surveys in biodiversity-rich, underexplored regions—especially targeting small-bodied, data-deficient taxa—and for timely updates and data-sharing to better inform conservation strategies.

## Introduction

Understanding species’ geographic distributions is fundamental to the fields of biogeography, macroecology, and conservation biology (Darwin, 1859, MacArthur, 1984, Whittaker et al. 2005, Diniz Filho et al. 2023), particularly in the face of the global biodiversity loss crisis. The status and trends of species’ spatial-temporal distributions are closely tied to species’ ecological functions, population dynamics, and extinction risks (Bland et al. 2015, Ribeiro et al. 2022), all of which are essential for effective biodiversity conservation and management (Pulliam, 2000, Guisan et al. 2013, Jetz et al. 2019, Oliver et al. 2021, Hughes et al. 2024). However, significant gaps and biases —referred to as the “Wallacean shortfall” —persist in our knowledge of species distributions, which describes the incomplete or inaccurate understanding of species’ spatial and temporal distribution ranges (Lomolino, 2004, Hortal et al. 2015), particularly in under-sampled areas and under-studied taxa (Guedes et al. 2024, Vieira et al. 2024). This shortfall undermines biodiversity assessments, skews ecological models, and may misdirect conservation actions (Lomolino, 2004, Hurlbert and Jetz 2007, Hortal et al. 2015, Hughes et al. 2021, 2024, Mentges et al. 2021, Diniz Filho et al. 2023, Bowler et al. 2024). Addressing these gaps is crucial for enhancing our understanding of biodiversity dynamics.

Species distributions are shaped by a complex interplay of intrinsic traits, environmental factors, biotic interactions and anthropogenic influences (Boulangeat et al. 2012, Di Marco et al. 2015, Pacifici et al. 2020, Comte et al. 2023, Feng et al. 2024). First, species traits, such as body size, habitat breadth, and dispersal capacity play a critical role in determining their distributions (Pacifici et al. 2020, Comte et al. 2023). For example, larger vertebrate species with broader habitat tolerances are often better able to adapt to changing conditions and colonize new environments (Calosi et al. 2010, MacLean and Beissinger 2017). Second, climate, topography, and habitat availability are pivotal environmental factors in shaping species’ ranges (Brown et al. 1996, Shuai et al. 2021), with climate change increasingly driving range shifts (Pecl et al. 2017, Pacifici et al. 2020, Schultz et al. 2022, Rubenstein et al. 2023). Third, biotic interactions (e.g., competition, predation) limit species’ ability to colonize or persist in certain habitats (Araújo and Luoto 2007, Wisz et al. 2013). Last, anthropogenic factors, such as urbanization and agriculture, further complicate species distribution patterns (Sirami et al. 2017, Pineda-Munoz et al. 2021).

Robust baseline data are essential for detecting global biodiversity changes and their impacts in a rapidly changing world (Pecl et al. 2017, Bonebrake et al. 2018, Oliver et al. 2021). Accurately documenting species’ distributions is not only a prerequisite for understanding their responses to environmental changes (e.g., climate change) but also for predicting future changes in biodiversity patterns and ecosystem functions (Elith et al. 2010, Pecl et al. 2017, Williams and Blois 2018, Pacific et al. 2020, Johnson et al. 2024, Lawlor et al. 2024). However, many previous studies have revealed significant taxonomic, spatial and temporal bias in species occurrence information, primarily due to uneven sampling efforts across space and time (Boakes et al. 2010, Beck et al. 2014, Meyer et al. 2015, 2016a, b, Hughes et al. 2021, Oliver et al. 2021, Bowler et al. 2022). Consequently, many biodiversity maps reflect sampling efforts more than actual species ranges (Hortal et al. 2007, 2015, Hughes et al. 2021). Specifically, geographical biases may result from biased fieldwork in terms of regional differences in socio-economic factors (Yang et al. 2014, Meyer et al. 2016b), accessibility (Monsarrat et al. 2019), funding (Wang et al. 2021), research capacity (Zhang et al. 2023), or a preference for areas rich in endemism, mountainous regions, or protected areas (Jetz et al. 2012, Meyer et al. 2015, 2016a, b). Additionally, biases towards certain species might stem from site-specific socio-economic factors (Meyer et al. 2016b) and/or from species-specific characteristics, such as the low detectability of nocturnal species (Zuberogoitia et al. 2020), arboreal species (Chen et al. 2023), cave-dweller species (Feijó et al. 2019), cryptic (Crawford et al. 2020) and threatened species (Boakes et al. 2010, Hu et al. 2019). These reflect the shortfall in monitoring, documenting, collecting and sharing biodiversity data and collectively emphasize the importance of data coverage, quality, and accessibility for effective biodiversity mapping and conservation efforts (Schmeller et al. 2017, Oliver et al. 2021, Bower et al. 2024).

Species inventory constitutes baseline data for the biota of a natural geographical region or administrative unit, and its establishing and timely updating holds significant importance for biodiversity conservation research and management (Ma, 2015, Jiang et al. 2022). In China, enhanced survey efforts, technological advances, and substantial investments have led to the establishment of multiple biodiversity monitoring platforms (Ma et al. 2018, Mi et al. 2021, Wu et al. 2022), such as China Biodiversity Observation and Research Network (Sino BON), China Biodiversity Observing Network (China BON). China has also built an openly accessible integrated bigdata platform for biodiversity and ecological security (BioONE, https://bio-one.org.cn/), as well as published a series of monographs and datasets on species inventory and distribution (e.g., Jiang et al. 2015, 2024), ecological traits (e.g., Wang et al. 2020, Ding et al. 2022), threatened status (e.g., Jiang et al. 2016, 2024). Despite these advancements, challenges such as lack of unified data collection standards, poor data integration, and limited data sharing hinder the effective use and deep exploration of biodiversity information (Zhang, 2017, Ma et al. 2018, Wei et a., 2021). For example, the coverage of China’s biodiversity data on globally accessible platforms such as GBIF is under 10% (Meyer et al. 2015, Oliver et al. 2021), and the data often suffer from spatial and temporal biases (Yang et al. 2014, Qian et al. 2018, Huang et al. 2020). This limited data availability and accessibility impedes a comprehensive understanding of China’s rich biodiversity (Huang et al. 2020, Mi et al. 2021, Wei et al. 2021)

Mammals are a relatively well-studied group among vertebrates (Burgin et al. 2018, Upham et al. 2019). China, as a mega-diverse country spanning parts of the Palearctic and Indomalayan realms, hosts 740 mammal species, accounting for 11.2% of global mammal species (Jiang et al. 2024). Of these, 182 species are endemic, highlighting the country’s unique mammalian fauna (Jiang et al. 2024). This diverse fauna includes nine of the world’s fourteen biomes, reflecting a high degree of heterogeneous climate, complex topography, and diverse ecosystems (Zhang, 2011, Jiang et al. 2015, Ding et al. 2022). Moreover, 174 and 94 species are assessed as ‘threatened’ and ‘data-deficient’, representing the red list status of China’s mammals (Jiang et al. 2024). While research on China’s mammals has advanced significantly in recent decades, driven by increased survey efforts and technological innovations (Huang et al. 2021, Mi et al. 2021), substantial gaps remain in the distribution data, particularly for elusive taxa such as bats (*Chiroptera*) and rodents (*Rodentia*) (Jiang et al. 2016, Chen et al. 2022). These groups are often overlooked potentially due to their small sizes, nocturnal behavior, and morphological similarities, making their detection and identification challenging.

In recent decades, Chinese zoologists have not only discovered many newly described mammal species, but also documented substantial new mammal distribution records (i.e., national or provincial-level newly recorded mammal species), which expand our understanding of mammalian ranges in China (Jiang et al. 2022, 2023, 2024). Such records not only alleviate the Wallacean shortfall but also help clarify species range boundaries, facilitate more accurate biogeographical delineations, and elucidate the processes that shape species distributions (Jiang et al. 2021, Ding et al. 2022). This study aims to investigate the spatiotemporal patterns and ecological correlates of new mammal records by analyzing data on national or provincial-level new mammal occurrences from 2001 to 2023 in China. Specifically, we seek to:

1. Characterize the taxonomic, spatial, and temporal patterns of new distribution records, hypothesizing that small body-sized, elusive taxa (e.g., Chiroptera, Eulipotyphla, Rodentia) that have historically received less research attention will dominate new records.
2. Investigate the drivers of new occurrence records by examining species traits (e.g., body size, habitat breadth), environmental conditions (e.g., climate, elevation), and socio-economic factors (e.g., GDP, human density) that influence the likelihood of discovering new mammal records. We also explore the relationships between provincial-level new records and species richness, historical (1949–2000) and current (2001–2023) survey efforts, as well as socio-economic indicators.
3. Identify major biogeographic trends and conservation implications, focusing on whether new occurrences represent the discoveries in previously unrecognized zoogeographical regions or expansions into novel zoogeographical regions or reflect disjunct distributions.

By integrating ecological, socio-economic and biogeographical data, this study provides a comprehensive framework for understanding the mammalian diversity and distribution in China. The findings will inform future biodiversity monitoring, guide conservation planning and resource allocation, contributing to efforts to bridge the critical knowledge gaps in biodiversity distribution research.

## 1.2. Material and methods

### 2.1 Data collection and handling

#### 2.1.1 New mammal occurrence data

We systematically collected new mammal occurrence records (excluding newly described species) from 2001 to 2023 in 34 provincial-level administrative regions (provinces, autonomous regions, and municipalities) across China. The data was sourced from the China National Knowledge Infrastructure (CNKI, https://www.cnki.net/) and Google Scholar (https://scholar.google.com). We retrieved all peer-reviewed publications using the following search terms: (China* OR Province) AND (mammal* OR mammalian OR wild mammals) AND (new occurrence records* OR new provincial-level records OR new distribution range OR new discovery OR new sighting OR new observation) (accessed in January 2024). After excluding records of rediscovery and taxonomic revisions without provincial-level distribution changes, we identified 192 papers representing 150 species in 26 families across 7 orders (Supporting information). From each paper, we extracted information on species identities, newly distributed provinces, locations of new occurrences, collection methods, potential reasons for occurrences and detections, habitat types, elevation, biomes, journal names, the years of new occurrences documented and published, and authors of the original papers. We then analyzed the spatio-temporal pattern of new mammal occurrence records.

#### 2.1.2 Explanatory variables for detections of new mammal occurrences

To identify potential predictor variables influencing the detections of new mammal occurrences, we compiled a comprehensive species list from the IUCN Red List (IUCN, 2024), which totals 640 mammal species. We used the species list from the IUCN Red List because it provides verified species names, standardized distribution maps, and threatened status information for extant mammals in China (Chen et al. 2023). We focused exclusively on terrestrial mammals, excluding 48 freshwater and marine species (Chen et al. 2023).

We categorized potential predictor variables into three groups: (1) Biological traits, including body size (mean adult body mass as a proxy), range size, activity breadth, habitat breadth, number of congeners, and the year of species description (Wilman et al. 2014, Cooke et al. 2019, Shuai et al. 2021, Ding et al. 2022, Chen et al. 2023). (2) Environmental conditions, such as mean temperature, mean precipitation, temperature seasonality, precipitation seasonality, and elevation (Moura and Jetz 2021, Shuai et al. 2021, Chen et al. 2023). (3) Socio-economic factors, including human density, GDP, and accessibility (i.e., travel time to high-density urban centers) (Weiss et al. 2018, Pouteau et al. 2022, Guedes et al. 2023, Mammola et al. 2023). See dataset S2 for more detailed information on the relationship between species attribute and new provincial-level distributions. Although research efforts (typically measured by the number of publications per species) affect detection likelihood (Zhang et al. 2023), we excluded it because most new records originated from broad biodiversity surveys targeting multiple taxa rather than species-specific studies (Wang et al. 2021, Jiang et al. 2022, Wu et al. 2023). Instead, we accounted for survey efforts by standardizing the number of publications per province associated with field surveys targeting mammals (Wu et al. 2023).

We extracted the species description date as the year when the species was formally described for the first time as an independent taxonomic unit (Diniz-Filho et al. 2005, Chen et al. 2023). We used mean adult body mass as a measure of body size for each species (Ding et al. 2021, Soria et al. 2021). Activity breadth was calculated as the number of foraging activity times (i.e., nocturnal, crepuscular, diurnal) (Ding et al. 2021). Habitat breadth was calculated as the number of suitable habitats per species according to the IUCN habitat classification scheme (Cooke et al. 2019, Jung et al. 2020). For species lacking relevant information on activity breadth (15 species) or habitat breadth (7 species), we imputed trait values by calculating the mean trait value from congeneric species in our dataset (Supporting information). The number of congeners was calculated as the number of mammal species in each genus in the year of the species’ formal description (Chen et al. 2023, Jiang et al. 2024). To obtain species’ geographic range size, we downloaded extent-of-occurrence maps from the IUCN Red List (accessed July 2024) and calculated range size in ArcGIS 10.8 (Esri, 2022). Mean elevation was derived from SRTM data (30 arc-second resolution) (https://www.worldclim.org/data/worldclim21.html<u>#</u>). We obtained human population density (Liu et al. 2024) and GDP data (Chen et al. 2022) at a 30 arc-second resolution for 2010, representing average human activity intensity and economic development levels during the study period, respectively. We also obtained accessibility data (Weiss et al. 2018, Nelson et al. 2019) to represent accessibility levels (Hickisch et al. 2019, Chen et al. 2023). Zoogeographical regions were assigned to each species following Gao et al. (2017a), delineating China into 2 realms, 7 regions, and 19 sub-regions based on Zhang (2011). Biomes were extracted from the Terrestrial Ecoregions of the World (https://ecoregions.appspot.com/) for each species (Olson et al. 2001, Ding et al. 2021). We estimated mean environmental conditions (i.e., climatic niche) by overlaying species range with bioclimatic variables (mean temperature, mean precipitation, temperature seasonality, precipitation seasonality) from 1970 to 2000 at a 30 arc-second resolution (WorldClim 2, Fick and Hijmans 2017).

To validate the environmental tolerance estimates for each species, we downloaded species occurrence data from the Global Biodiversity Information Facility (GBIF, https://www.gbif.org/) using the rgbif package (Chamberlain et al. 2021), and compiled occurrence data from the Chinese National Animal Collection Resource Center (NACRC, http://museum.ioz.ac.cn/), and Smith and Xie (2009). We cleaned these data using the CoordinateCleaner package (Zizka et al. 2019) by removing duplicate coordinates and problematic records (e.g., empty elements, non-georeferenced occurrences). Additionally, we excluded GBIF occurrences outside of IUCN polygons, retaining only points within each species’ IUCN range maps. For species with fewer than 30 occurrences, we used range maps to extract mean environmental values (Alhajeri and Fourcade 2019). We found no significant differences when using geographic range versus occurrence data in estimating species’ environmental tolerances (88.0-96.7% of species with geographic ranges fall within 95% confidence intervals with species occurrences in terms of bioclimatic value estimates, see dataset S3). Thus, we present results based on geographic distribution in the main text.

#### 2.1.3 Explanatory variables for the number of provincial-level new mammal occurrences

To explore factors influencing the number of new mammal records across provinces, we extracted the number of publications targeting mammal field surveys in China (1949-2023) from Wu et al. (2023) and Jiang et al. (2021, 2022, 2023). Then we divided these field surveys into two periods: 1949-2000 (historical) and 2001-2023 (current). We obtained mammal species richness for each province from Ding et al. (2022) and Jiang et al. (2024). We calculated the survey effort of each province as the number of publications divided by administrative area and species richness (Ding et al. 2021, Jiang et al. 2021, Wu et al. 2023). We compiled human density and per capita GDP data for each province from this platform (https://www.resdc.cn/).

### 2.2 Statistical analyses

To determine the direction of new mammal records relative to known distribution ranges before, we calculated the centroid of each species’ extent-of-occurrence polygon using ArcGIS 10.8. Then we computed the azimuth angle from the centroid to the new record. We classified directions into eight sectors: North, Northeast, East, Southeast, South, Southwest, West, and Northwest at 45-degree intervals.

To quantify the influence of species characteristics on new occurrence records (i.e., presence of new mammal records), we fitted Bayesian Phylogenetic Generalized Linear Mixed Models (BPGLMM) with a binomial error structure and a logit-link function, using Markov Chain Monte Carlo (MCMC) techniques in the MCMCglmm package (Hadfield, 2010) in R 4.4.3 (R Core Team, 2024). This approach accounts for phylogeny as a random effect (Pagel, 1999, Freckleton et al. 2015). Considering flying bat species may exhibit distinct distributional patterns compared to non-flying mammals, we fitted Bayesian models for all mammals, non-flying mammals, and bat taxa separately. To control for non-independence due to shared ancestry, we included a phylogenetic covariance matrix as a random effect (Freckleton et al. 2002). The phylogenetic variance–covariance matrix was generated using the vcv function in the phytools package (Revell, 2012), which contains pairwise shared evolutionary history based on branch lengths of the phylogenetic trees (He et al. 2023). The mammalian phylogenetic tree was constructed as a maximum clade credibility tree using BEAST, based on 5,911 mammal species from 1,000 randomly sampled trees from Vertlife (Upham et al. 2019). We excluded 15 species lacking phylogenetic information. As a result, we compiled 534 terrestrial mammal species: 187 Rodentia, 127 Chiroptera, 74 Eulipotyphla, 45 Cetartiodactyla, 39 Carnivora, 31 Lagomorpha, 25 Primates, 4 Perissodactyla, 1 Pholidota, and 1 Scandentia. All data used in this study are provided in supporting information.

To account for phylogenetic uncertainty, we performed the models across a random sample of 1,000 phylogenetic trees, summarizing slope estimates using mean values. We ran two MCMC chains for 55,000 iterations each, discarding the first 5,000 as burn-in and using a thinning interval of 50, yielding an effective sample size of 1,000. We verified that autocorrelations were below 0.1 and that chain convergence was below 1.1. We specified weakly informative priors (V = 1, nu = 0.002) for phylogenetic and residual variances, following Hadfield (2010). We performed posterior predictive checks to validate the model by comparing simulated values with observed data using the pp_check function in bayesplot (Gabry and Mahr 2022). We reported the medians and 95% credible intervals of all parameter estimates from the posterior distributions and considered the significance of the estimate if the 95% credible interval for the coefficient did not overlay with zero (He et al. 2023, Zhong et al. 2024). In the full model, we modeled newly recorded mammal species as a function of body size, activity breadth, habitat breadth, number of congeners, range size, temperature seasonality, precipitation seasonality, mean elevation, elevation range, and human density. Before formal analyses, all variables were log10-transformed and standardized (mean = 0, SD = 1). We assessed multicollinearity via the Variance Inflation Factor (VIF) in the car package. Since all VIF < 5, we retained all predictors.

To analyze the drivers of provincial-level new occurrence records, we constructed a Multivariate Poisson Regression Model, with the number of new records per province as the response variable, and human density, per capita GDP, mammal species richness, and historical survey efforts (1949-2000) and current survey efforts (2001-2023) as predictors. We then performed partial regression analyses to quantify relationships between the number of newly recorded mammals in each province and each predictor, controlling for the others (Carrascal et al. 2009). Given potential multicollinearity (Chen et al. 2023), we used hierarchical partitioning to assess the relative importance of each variable (Lai et al. 2022). Specifically, hierarchical partitioning calculates and partitions goodness-of-fit for all potential models, summing each predictor’s unique and average shared variance. This method enables the interpretation of colinear variables’ contributions, whereas forward or backward stepwise approaches cannot. We performed hierarchical partitioning using the glmm.hp (0.1-5) package, determining both shared and individual variance for the new occurrence records across all possible model combinations (Lai et al. 2022).

## 3. Results

### 3.1 Distribution of new mammal records in China

From 2001 to 2023, 150 new provincial-level mammal records were documented in China (Table 1), encompassing 8 orders and 26 families. Chiroptera (bats) accounted for the highest number of the new records (n = 69, 46%), with 46 species belonging to Vespertilionidae (Vesper bats) and 13 species to Rhinolophidae (horseshoe bats). Additionally, Hipposideridae (Old World leaf-nosed bats) contributed 5 new records, representing 55.5% of their total family species counts. Eulipotyphla significantly contributed 26 new records, with 23 species from Soricidae. Rodentia added 23 new records, including 8 species from Muridae and 7 species from Cricetidae. Of these, 42 (28%), 12(8%) mammal species are listed as “data deficient” and “threatened” status in the IUCN Red List, respectively (Fig. 1).

**Fig. 1.**
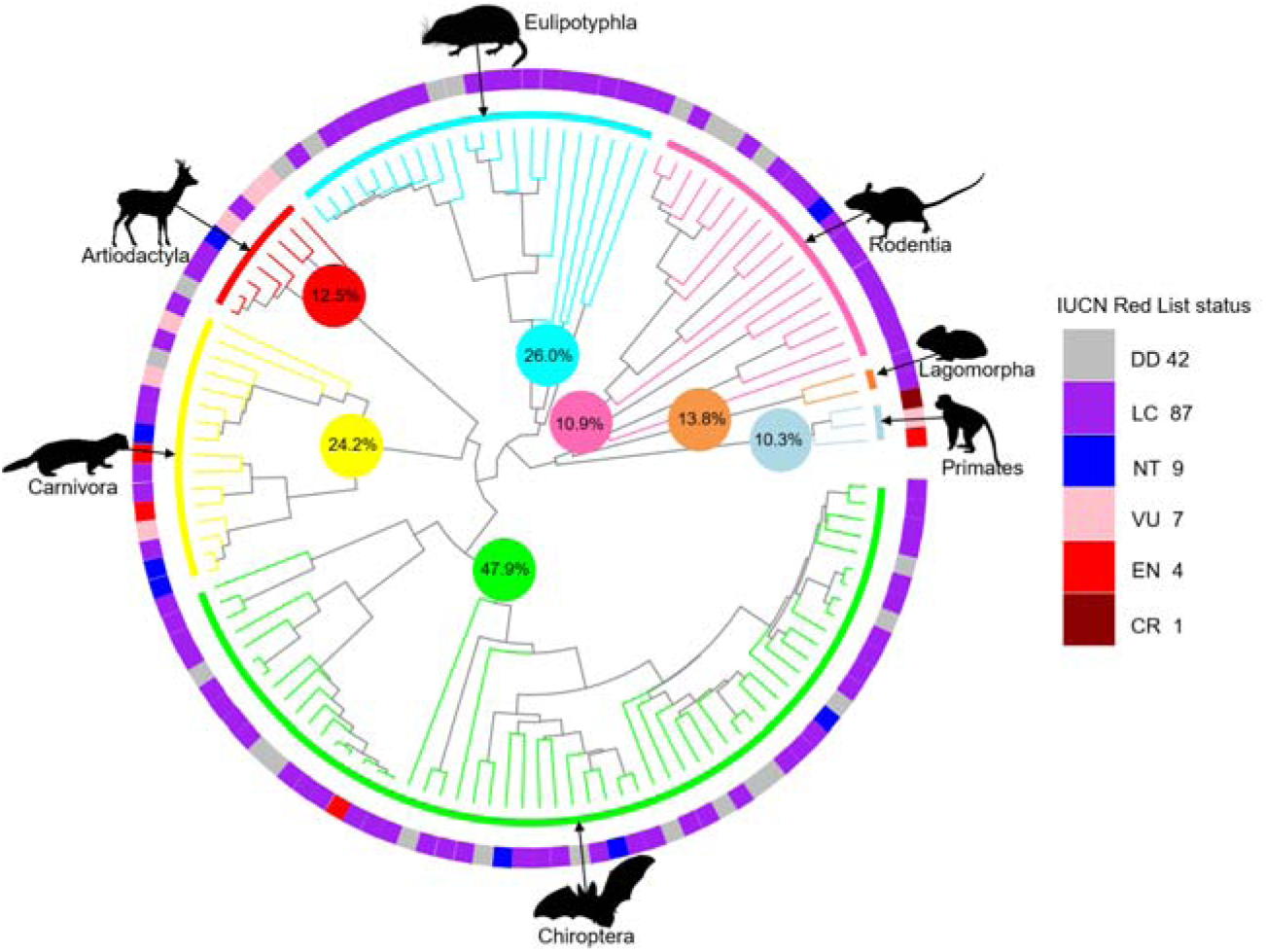
The phylogenetic tree illustrates newly recorded mammal species in China between 2001 and 2023, categorized by taxonomic order and IUCN Red List conservation status. This phylogeny was generated using the R package “V.PhyloMaker” (Jin and Qian 2019) based on the backbone of phylogenies from Uphamet al. (2019) and was randomly drawn from 1,000 phylogenetic trees. The outermost circular band of the tree shows IUCN red list status of each newly recorded mammal species, while the inner colored-rings and phylogenetic branches indicate the different taxonomic orders, and percentages inside colored circles represent the proportion of new mammal records to total number of species of the order in China. Each number next to the IUCN red list status indicates the number of newly recorded species. Animal silhouettes for representative species in orders are shown and are downloaded from phylopic.org under public domain license.

**Fig. 2.**
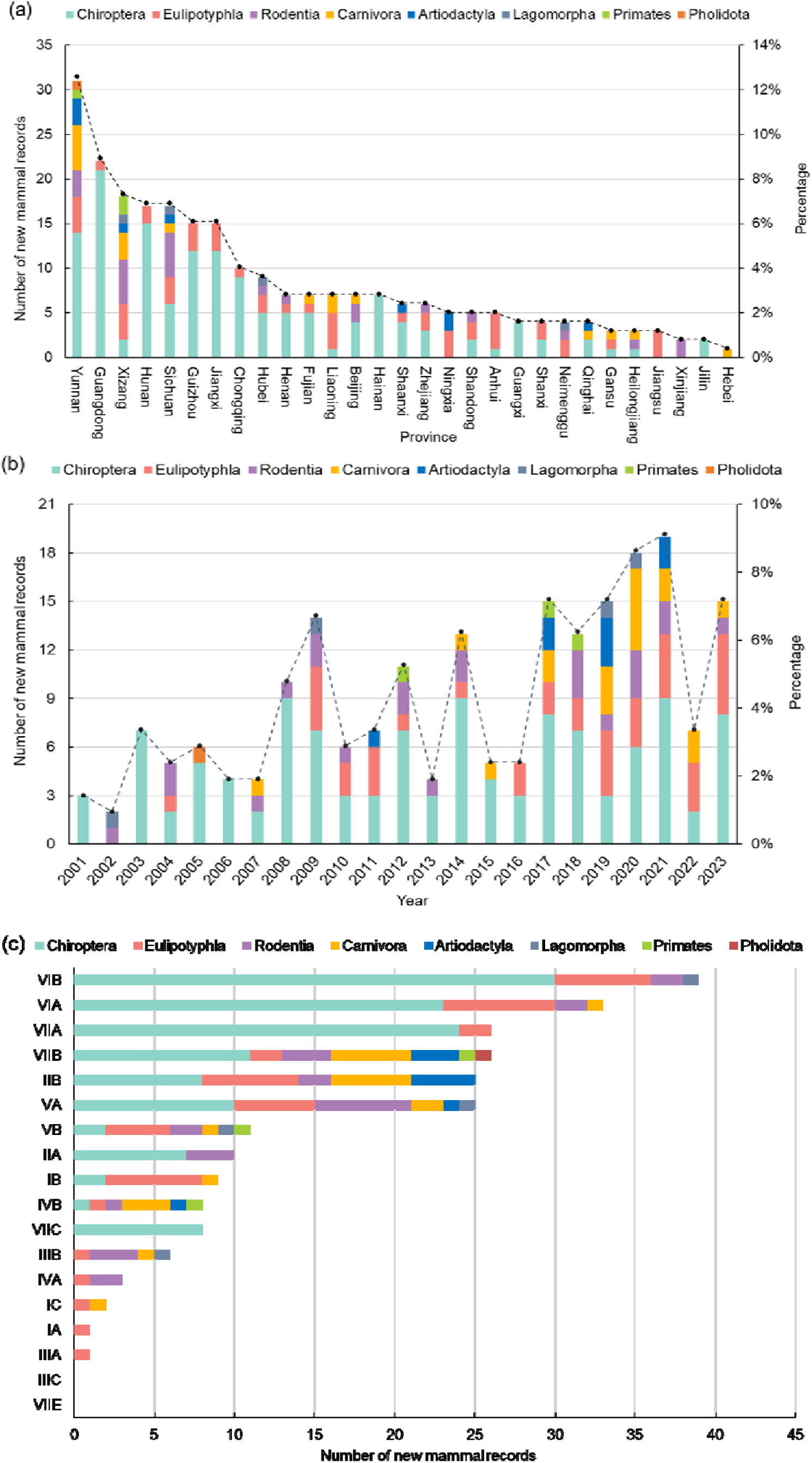
Number of new mammal records across provinces (a) and per year (b) and distribution of newly recorded mammal species across zoogeographic subregions, categorized by taxonomic orders in China between 2001 and 2023. Note: Zoogeographic subregions are classified based on geographical features of species distribution (see Zhang, 2011 and Gao et al. 2021), and bars represent the number of new mammal species recorded for each order. IA: Daxing’anling Mountain subregion, IB: Changbaishan Mountain subregion, IC: Song-liao Plain subregion, IIA: Huang-huai Plain subregion, IIB: Loess Plateau subregion, IIIA: East Steppe subregion, IIIB: West Desert subregion, IIIC: Tianshan Mountain subregion, IVA: Qinghai-South Xizang subregion, VA: Southwest Mountain subregion, VB: Himalaya subregion, VIA: (East Hilly Plain subregion, VIB: Western Mountains and Plateaus subregion, VIIA: Min-Guang Coast subregion, VIIB: South Yunnan Mountain subregion, VIIC: Hainan subregion, VIID: Taiwan subregion, VIIE: South China Sea Islands subregion.

**Table 1.**
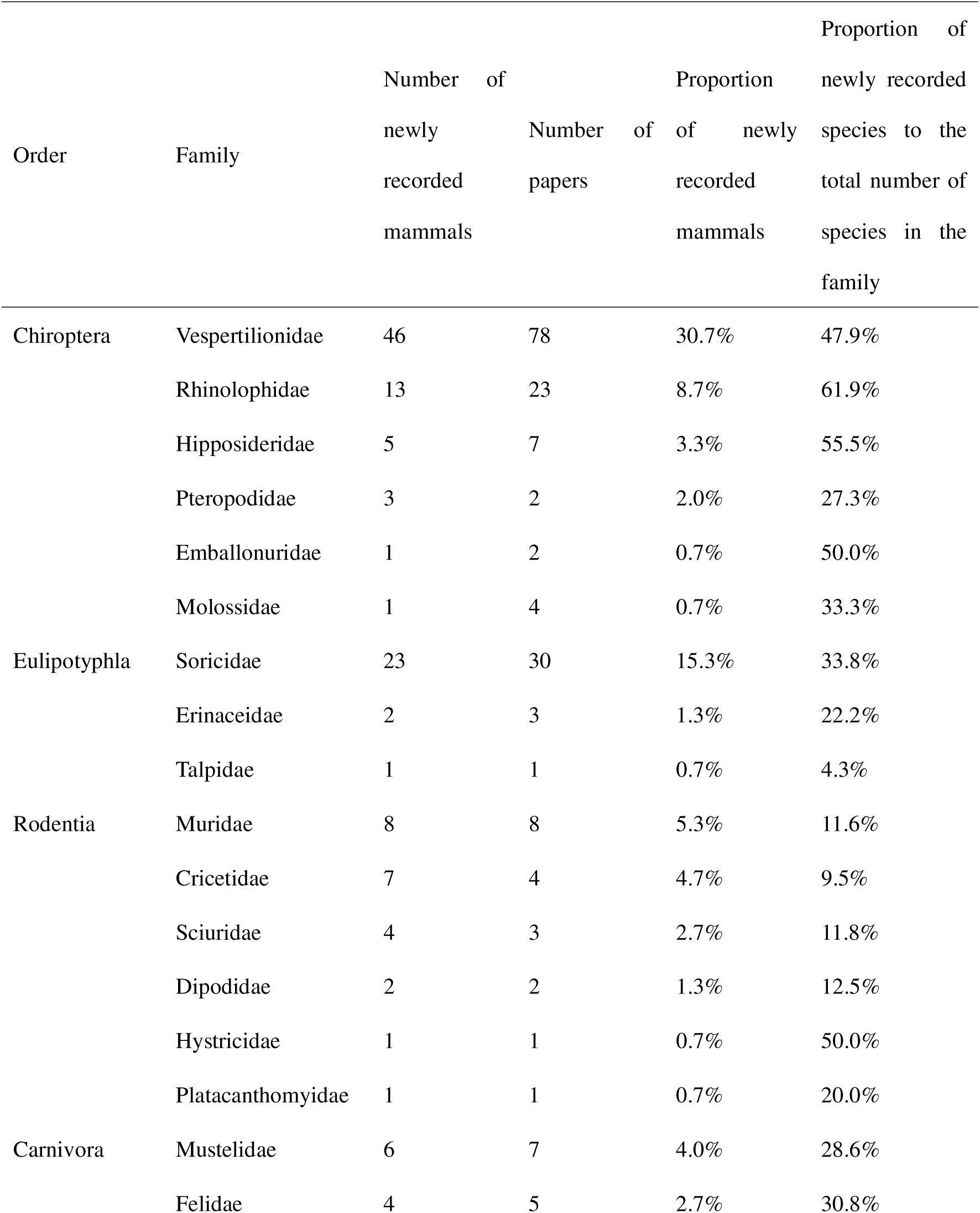
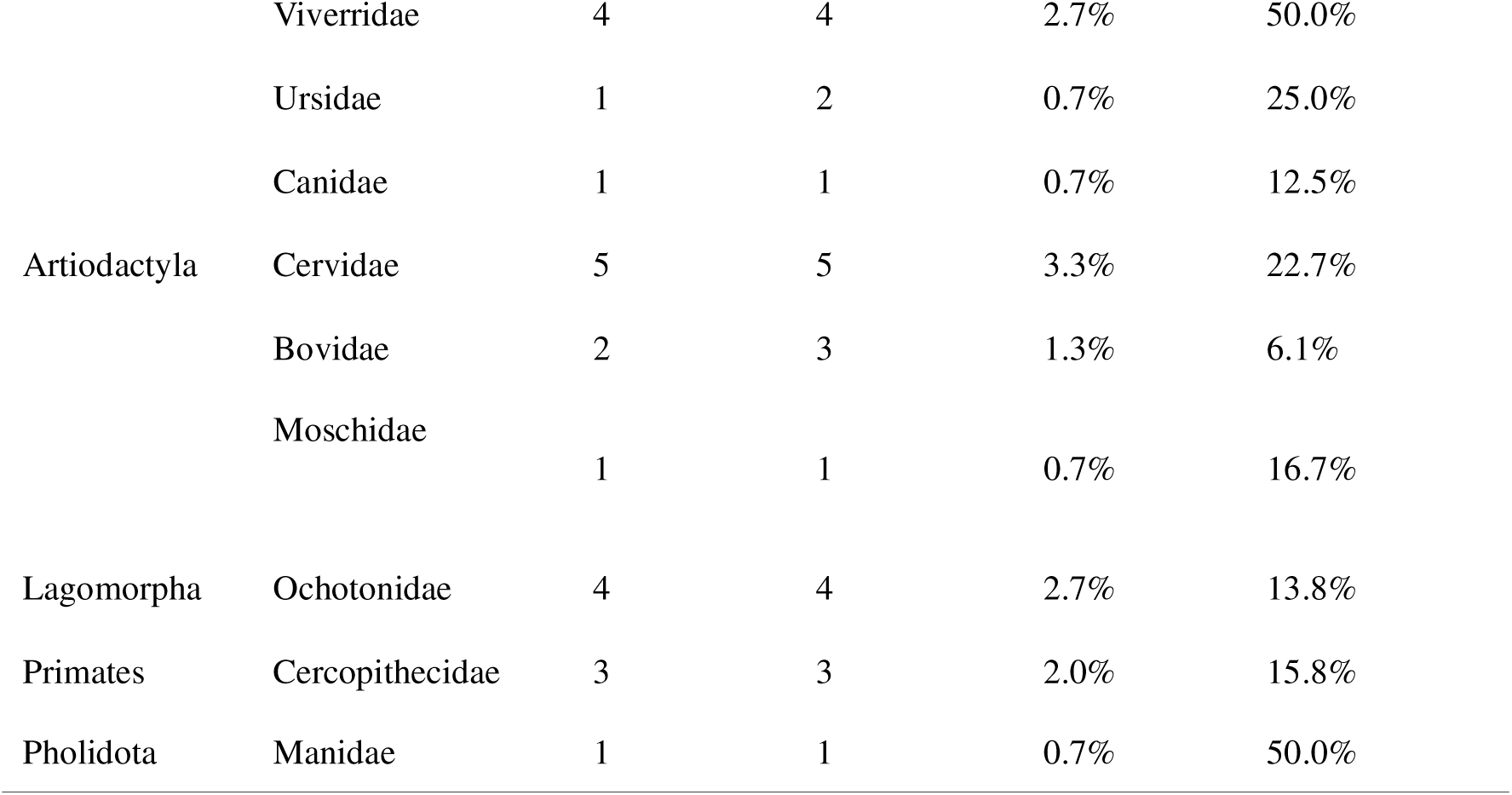
Summary statistics of newly recorded mammal species in China between 2001 and 2023

Geographically, the southern provinces—Yunnan (n = 31), Guangdong (n = 22), and Xizang (n = 18)—recorded the highest number of new mammal records over the past two decades (Fig. 1a, Fig S1). In contrast, northern provinces such as Xinjiang, Jilin, and Hebei reported fewer new records, averaging fewer than five discoveries each. Chiroptera dominated most provinces, particularly in Yunnan, Guangdong, and Sichuan. Rodentia and Eulipotyphla also made substantial contributions, especially in Yunnan, Sichuan, and Xizang. Annually, the number of new mammal records fluctuated, with notable peaks between 2017 and 2021 (Fig. 1b). The median time between discovery and publication of new records is 19 months (See supporting information dataset S2). Nineteen new records were documented in 2021. Chiroptera led in all years except 2002, while Rodentia and Eulipotyphla showed significant contributions in 2009, 2017, and 2021. Other orders, such as Primates and Carnivora, were sporadically represented.

At the zoogeographical subregional level, the Western Mountain and Plateau (VIB) subregion recorded the highest number of new species (n = 39), dominated by Chiroptera (n = 30) (Fig. 1c, Fig. S2). The East Hilly Plain (VIA) subregion followed with 33 new species, featuring a balanced representation of Chiroptera and Eulipotyphla. Chiroptera were particularly prominent in southern subregions such as South Yunnan Mountain (VIIB) and Hainan (VIIC). In contrast, subregions like the Loess Plateau (IIB) and Qiang-Tang Plateau (IIIC) reported fewer new records. Notably, 66 species were found in new zoogeographical subregions, 20 species occurred in discontinuous zoogeographical subregions relative to historically known distribution ranges (i.e., disjunctive distribution). Of these, two species—*Elaphodus cephalophus* and *Muntiacus reevesi*— both endemic to China and historically considered to be Oriental species, recorded in the Palearctic realm and occurred in the Liupan Mountain National Nature Reserve, Ningxia Hui Autonomous Region.

### 3.2 Direction of newly recorded species relative to their original distribution

Newly recorded species predominantly occurred towards the north and east relative to their known ranges, especially among small, cryptic mammals like bats and shrews (Fig. 4, Fig. S3). Chiroptera dominated in nearly all directions, contributing the highest proportion of new records in the North (n = 42, 35.0%), East (n = 31, 25.8%), and Northeast (n = 15, 12.5%). Eulipotyphla were prominent in the East (n= 10, 22.7%) and West (n = 10, 22.7%), whereas Rodentia in the North (n = 9, 33.3%) and Northwest (n = 8, 29.6%). Carnivora, Artiodactyla and Primates showed high proportion of new records in the northern and northwestern directions.

**Fig. 4.**
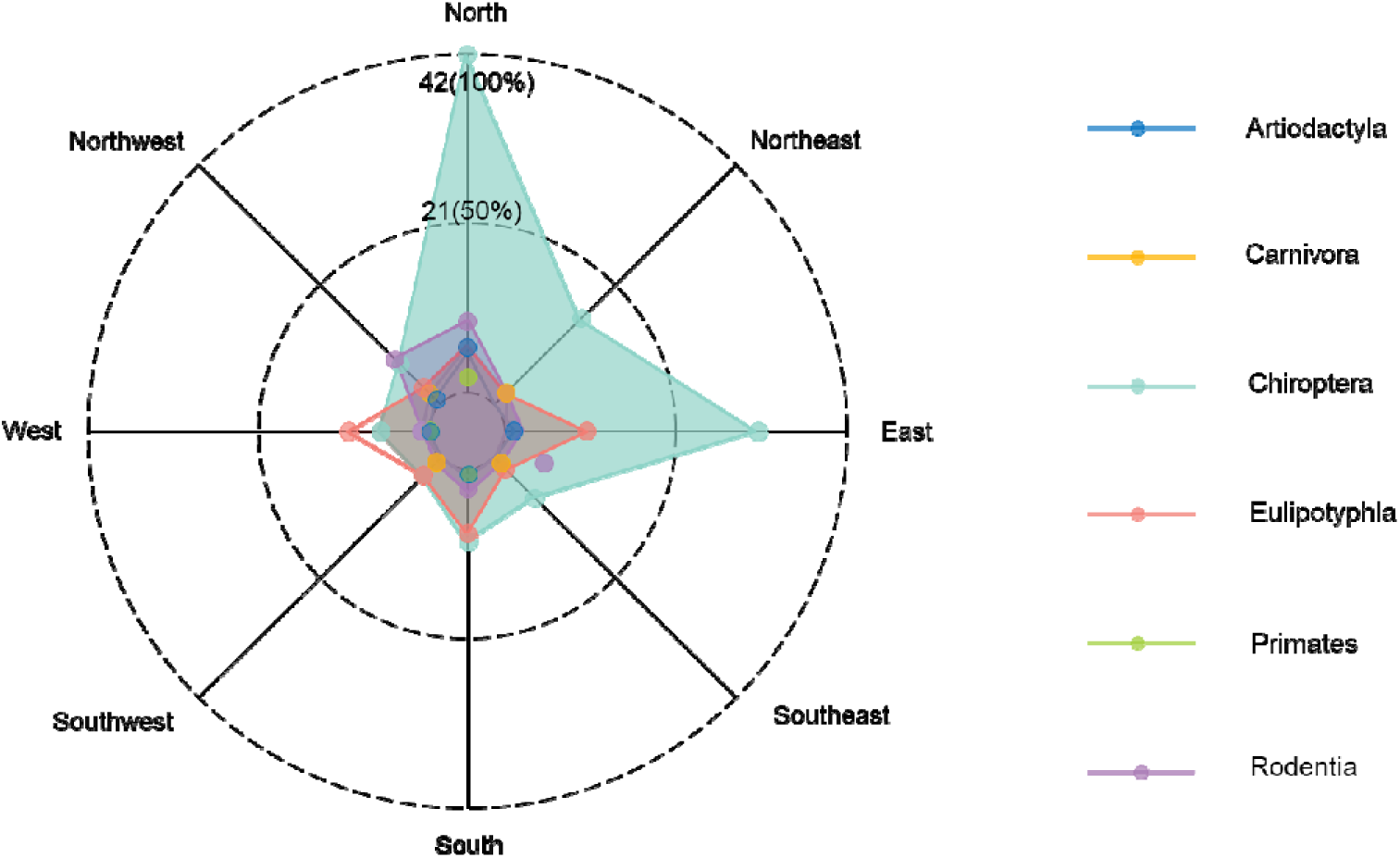
The directions of new mammal records occurred relative to their known distribution ranges. The numbers and the proportions in brackets marked on the north line represent the number and proportion of newly recorded species in Chiroptera, respectively.

### 3.3 Reasons and methods for discovering new mammal records

The majority of new mammal records (89.3%) were attributed to incomplete surveys and insufficient research in the past (Fig. S4). A smaller proportion (4.2%) was linked to range expansions or shifts, potentially due to climate change, habitat alterations, or human influence. Other reasons included species reclassification and advancements in taxonomic methods.

In terms of methodologies, traditional fieldwork was the primary method for identifying new records, accounting for 69.6% of discoveries. This involved the collection of specimens from the wild. Species identified with noninvasive camera-trapping techniques contributed 10.7%, particularly for elusive or nocturnal species. Species identified by re-examining historical museum specimen collections added 2.2% to new records. Other sources—including the species identified from those obtained by law enforcement, direct observations, and incidental captures—accounted for 17.4% of the new discoveries.

### 3.4 Relationships between new mammal records and predictor variables

The Bayesian PGLMM analysis revealed that body size exhibited a significant negative effect on the likelihood of new records for all mammal groups and non-flying mammals (Fig. 5). In contrast, habitat breadth and accessibility showed a consistent positive relationship with these groups. However, these effects were less pronounced for bats. The year and activity breadth variables consistently showed positive but statistically non-significant effects on all groups. Human density and GDP showed positive but statistically non-significant effects on all mammals and non-flying mammals, whereas the negative relationship was with bats. Range size emerged as a strong positive predictor for bats, while accessibility exhibited a statistically meaningful positive relationship for non-flying mammals. Elevation had a significant positive effect on non-flying mammals, while its influence was less clear for bat groups. The number of genera did not significantly influence discoveries.

**Fig. 5.**
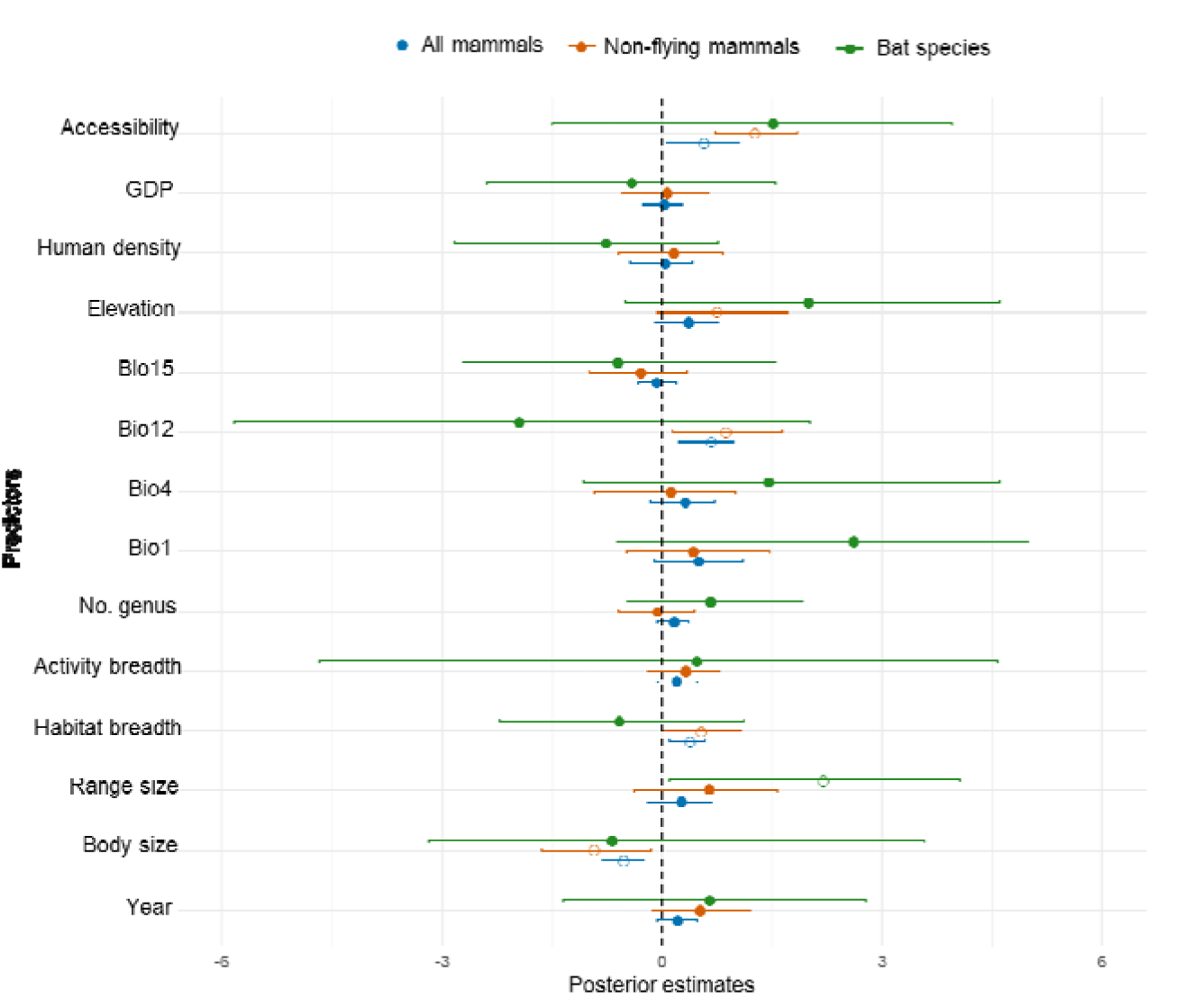
Distributions of the posterior estimates from Bayesian Phylogenetically Controlled Generalized Linear Mixed Models for predictors of new distribution records across all mammals (blue), non-flying mammals (orange), and bat species (green). The dots and error bars represent the median values and 95% posterior credible intervals, respectively. Unfilled dots indicate 95% credible interval for the posterior estimates did not overlap with zero (pMCMC < 0.05). The dashed line showing the location of the posterior mean slope estimate equals zero.

In addition, annual precipitation (Bio12) was an important predictor for non-flying mammals, showing a notable positive effect. In contrast, annual mean temperature (Bio1), temperature seasonality (Bio4) and precipitation seasonality (Bio15) did not show significant association with the likelihood of new mammal records. Overall, smaller-bodied species and those with broader habitat tolerances were more frequently discovered.

### 3.5 Relationships between provincial-level number of new mammal records and predictor variables

The multivariate Poisson regression analysis (Fig. 6) revealed that current survey efforts had a strong positive correlation with the number of new occurrence records (R = 0.78, p = 0.044), indicating that more active research leads to increased species discoveries. In contrast, historical survey efforts (1949-2000) showed a weak negative correlation (R = -0.42, p = 0.071), suggesting diminishing returns in areas that were already well-studied. Species richness was positively correlated with new species discoveries (R = 0.72, p = 0.032), indicating that biodiversity-rich areas tend to yield more new records. Per capita GDP exhibited a moderate positive correlation (R = 0.45, p = 0.051), suggesting that wealthier regions may support more research activities, indirectly enhancing species discovery rates. Hierarchical partitioning indicated that per capita GDP and species richness were the most influential predictors of provincial-level new mammal records, accounting for the majority of the explained variance, while historical survey efforts and contributed less significantly (Fig. 6).

**Fig. 6.**
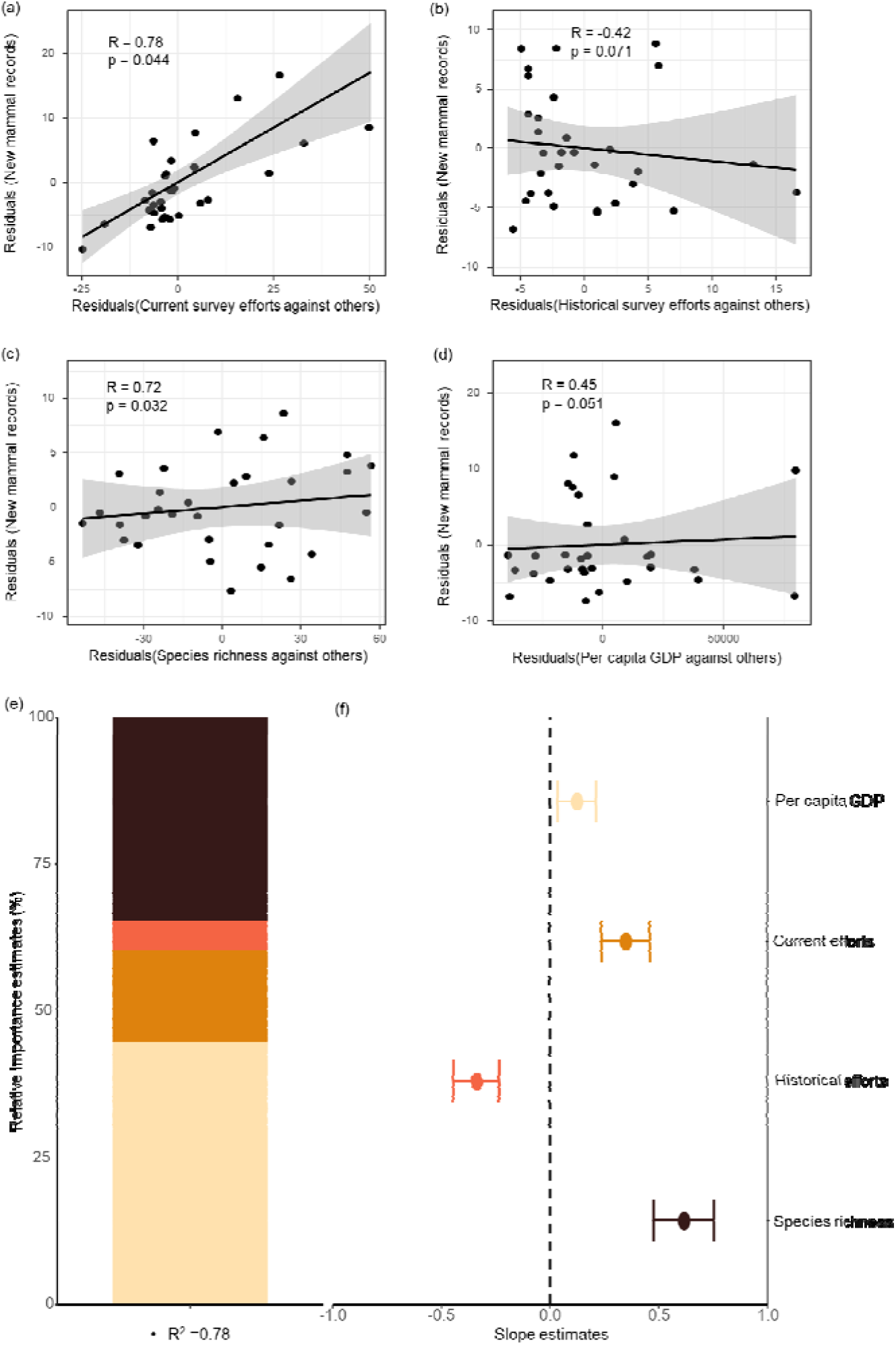
Partial regression plots and relative importance estimates showing the relationships between the residuals of newly recorded mammal species and key predictors. The predictors include current survey efforts (a), historical survey efforts (b), species richness (c), and Per capita GDP (d). Each plot displays the residuals of the newly recorded mammals regressed against each predictor while holding other variables constant. The lines represent the fitted regression models with 95% confidence intervals, along with the correlation coefficients (R) and p-values. The relative importance estimates (e) and slope estimates (f) of predictor factors influencing the number of provincial-level new mammal records. Note: the error bars with different colors correspond to the predictor factors labeled on the y axis (e), and this color scheme also applied in the colored column chart (f).

## 4. Discussion

The discovery of 150 new mammal distribution records across China from 2001 to 2023 highlights the importance of systematic biodiversity surveys in addressing the Wallacean shortfall and advancing conservation biogeography. These findings underscore the persistent gaps in mammalian distribution data. By synthesizing data on new distribution records, species traits, and anthropogenic factors, this research not only enriches our understanding of China’s mammalian distribution but also provides foundational data for species conservation and insights for prioritizing research efforts.

### 4.1 Taxonomic patterns and sampling biases

The dominance of Chiroptera, Eulipotyphla and Rodentia among the newly recorded species aligns with global trends where smaller, elusive taxa often remain underrepresented in biodiversity surveys and global research observations, as well as conservation efforts (Hu et al. 2019, Meyer et al. 2015, Hortal et al. 2015). The characteristics of these groups, such as small body size, specialized habitats, and cryptic behavior, pose challenges to detection and identification (Feijó et al. 2019, Zuberogoitia et al. 2020). This taxonomic bias highlights the critical need for targeted survey methodologies, such as eDNA techniques, DNA barcoding and bioacoustic monitoring, which have shown promise in detecting cryptic and elusive species (Crawford et al. 2020, Jiang et al. 2022). The reliance on traditional field methods for 69.6% of the new records emphasizes the continued importance of direct specimen collection in biodiversity research. However, the increasing role of non-invasive methods, such as camera traps and acoustic sensors, indicates a paradigm shift in survey strategies, enabling broader taxonomic and geographical coverage while minimizing ecological impact (Chen et al. 2023). Furthermore, improving survey and research efforts for the diverse but understudied taxa will mitigate the Wallacean shortfall (Haelewaters et al. 2024). Future studies should integrate these technologies with citizen science initiatives to enhance sampling coverage and research intensity in underexplored regions and understudied taxa (Bowler et al. 2024).

### 4.2 Spatial and temporal dynamics of new records

The spatial concentration of new records in southern provinces, such as Yunnan, Guangdong, and Xizang, reflects their high species richness and heterogeneous landscapes, which provide diverse habitats conducive to mammalian diversity (Zhang, 2011, Jiang et al. 2024). These regions also benefit from relatively intensive survey efforts and investments in biodiversity monitoring platforms, such as the China Biodiversity Observation Network for Mammals (China BON-mammal, http://114.251.10.194/html5/mammal/) (Wan et al. 2020, Guo et al. 2022) and Sino BON Mammal Diversity Monitoring Network (Sino BON- Mammal) (Xiao et al. 2023). Conversely, the paucity of new records in northern provinces highlights the limitations of historical survey efforts and lower mammalian diversity in these areas. The temporal peaks in new discoveries during 2017–2021 are attributed to increased funding, technological advances, and heightened research interest following the implementation of national biodiversity initiatives (Huang et al. 2021). However, the diminishing returns in historically well-surveyed regions underscore the need to prioritize underexplored areas to maximize the efficiency of conservation resources (Beck et al. 2014).

Previous studies have shown that many birds in China have shifted their distributions northward in response to climate change (Wu and Zhang 2015, Yang et al. 2020, Xing et al. 2024). However, the observed northward and eastward range shifts of newly recorded mammals should mainly be attributed to sampling bias rather than climate-induced range shifts. However, climate-induced range expansions or contractions have been substantially recorded (Chen et al. 2011, Pecl et al. 2017, Pacifici et al. 2020, Rubenstein et al. 2023). For example, Asian house rats have shown a noticeable range expansion from the southeast (warm and wet) to the northwest (cold and dry) of China since 1980, associated with an increase in air temperature and human activities (Bai et al. 2023). These shifts emphasize the need to integrate climate change models into biodiversity monitoring frameworks to predict and mitigate future distributional changes, as well as suggest that future research should focus more on the distribution dynamics of marginal populations (e.g., leading edge and trailing edge of species’ distribution) in the face of rapid climate and land-use changes (Hill et al. 2011).

### 4.3 Ecological correlates and drivers of new mammal records

The significant influence of body size on new mammal records aligns with historical studies demonstrating that smaller-bodied species are more likely to lack research attention and evade detection until recent targeted surveys (MacLean and Beissinger 2017, Pacifici et al. 2020). Specially, although smaller species are often more abundant and have larger population sizes, making them more likely to be found, they may have been historically overlooked due to sampling biases, as little research attention has been paid compared to large species and traditional survey methods may have been less effective in detecting them (Shuai et al. 2021, Chen et al. 2023). Species with wider habitat tolerances are more adaptable and occupy a greater variety of ecological niches, increasing their chances of being detected in different regions.

The strong positive correlation between current survey efforts and new records indicates that increased investment in biodiversity research, such as more extensive field surveys and advanced survey techniques, is crucial for uncovering historically unknown species distributions. This finding is consistent with previous studies that have emphasized the importance of research attention and funding in documenting biodiversity and promoting species conservation (Hu et al. 2019, Wang et al. 2021, Wu et al. 2023). Therefore, future research should focus on less explored regions or taxa to maximize the discovery of new species and new distribution records (Moura and Jetz 2021). Additionally, the positive association of GDP and human density with new records highlights the indirect role of socio-economic development in facilitating research activities and improving accessibility (Monsarrat et al. 2019, Pouteau et al. 2022), suggesting that conservation efforts in more developed regions may indirectly benefit from higher research activity and species discovery.

### 4.4 Implications for Conservation Biogeography

Extending known ranges into previously unrecognized zoogeographical realms and subregions has significant implications for biogeographical delineations and conservation planning. For instance, the presence of *Elaphodus cephalophus* and *Muntiacus reevesi* in the Liupanshan Nature Reserve of Ningxia, is the northern edge of their known ranges (Gao et al. 2017b, Luo et al. 2019). These discoveries in the Palearctic realm challenge traditional zoogeographical boundaries and highlight the dynamic nature of species distributions under changing environmental conditions (Gao et al. 2017a, Jiang et al. 2021). More research attention should be paid to species range boundary (Thomas, 2010). Additionally, the capture of a golden cat (*Catopuma temminckii*) at an altitude of 4,415 m in Medog County, Xizang, updates the upper altitude limit of this species’ distribution (https://www.news.cn/local/20240126/6a8615b78fae4f968e768ddf730412e0/c.html). The finding underscores the importance of maintaining habitat connectivity to facilitate species dispersal and resilience to environmental changes. Protected areas in biodiversity-rich regions, such as Yunnan and Xizang, should be prioritized for conservation investments to safeguard critical habitats and support species with shifting ranges (Jiang et al. 2016, Mi et al. 2021). This study also emphasizes the need for targeted conservation efforts in biodiversity-rich regions and for data-deficient taxa.

### 4.5 Addressing the Wallacean Shortfall

Despite significant advancements, substantial gaps remain in mammalian distribution data, particularly for data-deficient and cryptic taxa. Standardizing data collection methodologies and fostering collaborations among research institutions will improve data quality and accessibility (Schmeller et al. 2017). Furthermore, enhancing data-sharing mechanisms, such as integrating provincial-level records into global databases like GBIF and IUCN, is essential for reducing spatial and temporal biases (Meyer et al. 2015, Oliver et al. 2021). Every new distribution record increases our knowledge of species ranges and is likely contributing to mitigating the Wallacean shortfall. Future studies should focus on: 1) Underexplored taxa and regions: targeted surveys for cryptic and understudied species in northern and arid regions; 2) Advanced detection methods: employing acoustic sensors, environmental DNA (eDNA), and artificial intelligence (AI) for species detection and identification (Lahoz-Monfort and Magrath 2021); 3) Long-term monitoring: establishing longitudinal studies to assess the impacts of climate change and human activities on mammalian distributions, 4) Integrative approaches: combining ecological, phylogenetic, and socio-economic data to develop predictive models for species distributions and conservation priorities.

## Conclusions

Our study synthesized and analyzed data on new mammal occurrence records in China, revealing spatiotemporal patterns and ecological correlates that provide novel insights into the distribution of mammal species. The discovery of 150 new records, predominantly among Chiroptera, Eulipotyphla and Rodentia, highlights the persistent gaps in mammalian distribution data and underscores the importance of targeted biodiversity surveys. Our findings indicate that small-bodied species with broader habitat tolerances are more likely to yield new records and that regions with higher survey efforts and greater species richness are hotspots for species discoveries. To address the Wallacean shortfall, it is essential to conduct long-term monitoring and targeted surveys. Additionally, integrating multiple data sources and enhancing data-sharing mechanisms will improve the comprehensiveness and accuracy of biodiversity databases. Moreover, developing species-specific conservation strategies that consider the adaptability and resilience of species under climate change and human disturbances is crucial. By compiling the new occurrence data, this study also provides a foundation for future research and contributes to the global effort to understand species distribution and conserve biodiversity in the face of rapid environmental change.

## Supporting Information

**Fig. S1.**
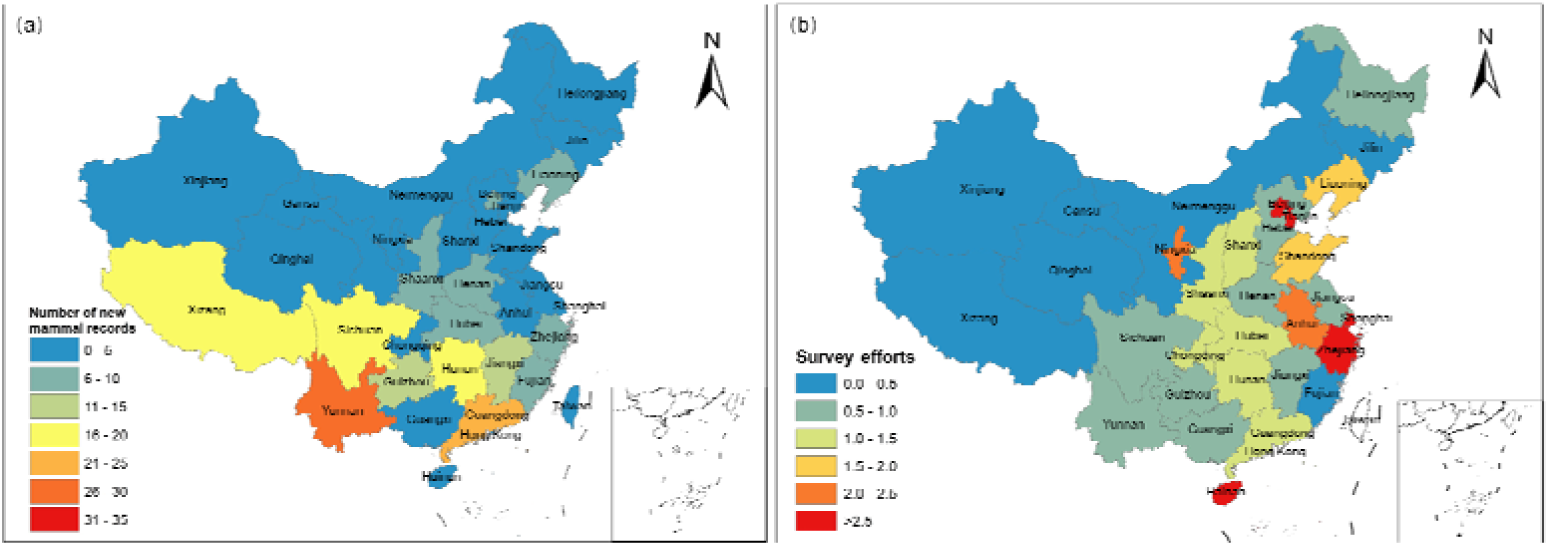
The spatial patterns of provincial-level new mammal records (a) and survey efforts (b) between 2001 and 2023 in China. Note: the survey effort was calculated as the number of publications targeted for mammal field surveys of each province divided by administrative area and mammal species richness.

**Fig. S2.**
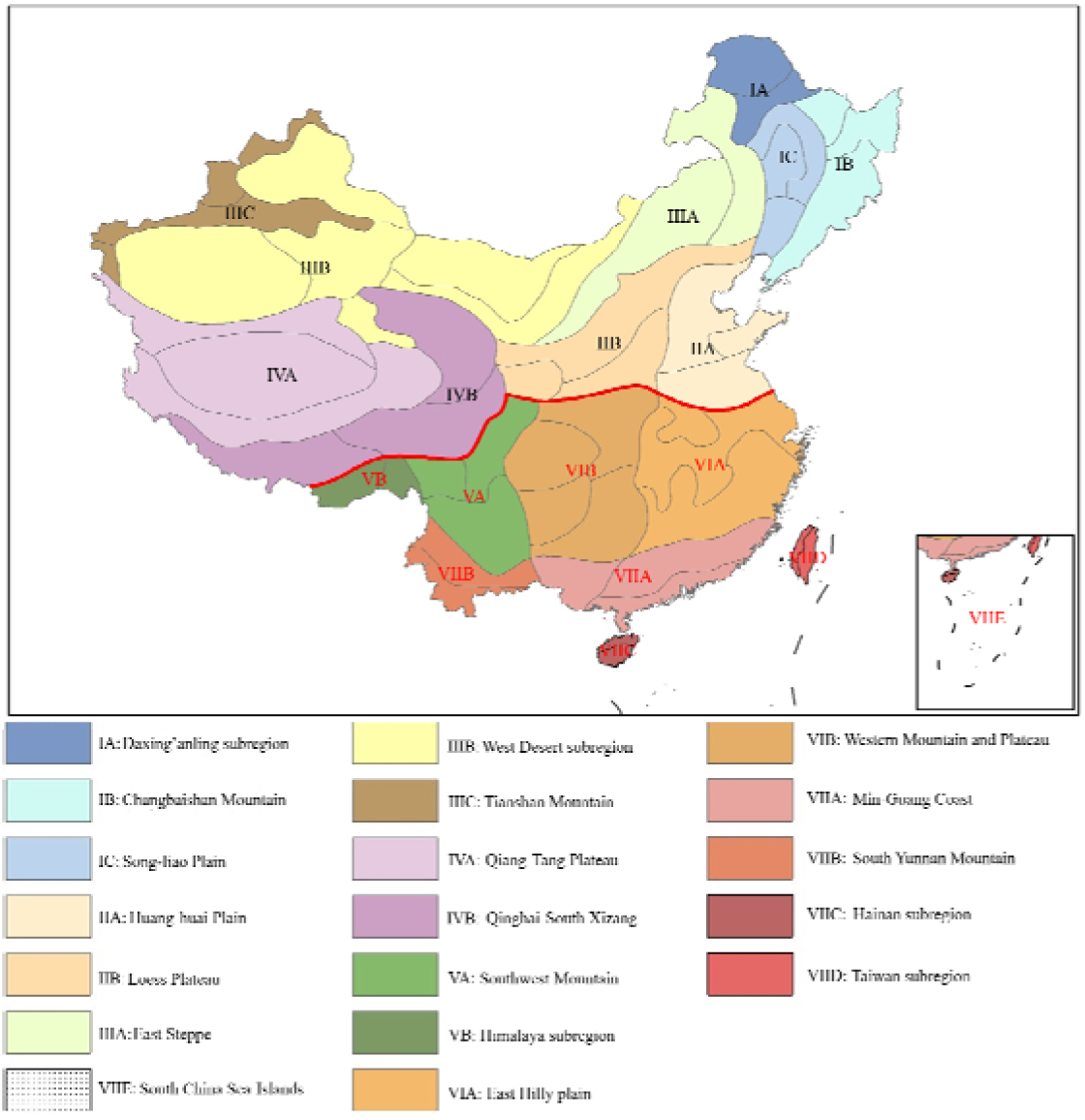
Map of zoogeographical sub-regions of China for terrestrial vertebrates (Adapted from Gao et al. 2017). The solid red line across it represents the boundary between the Palearctic and Oriental realms.

**Fig. S3.**
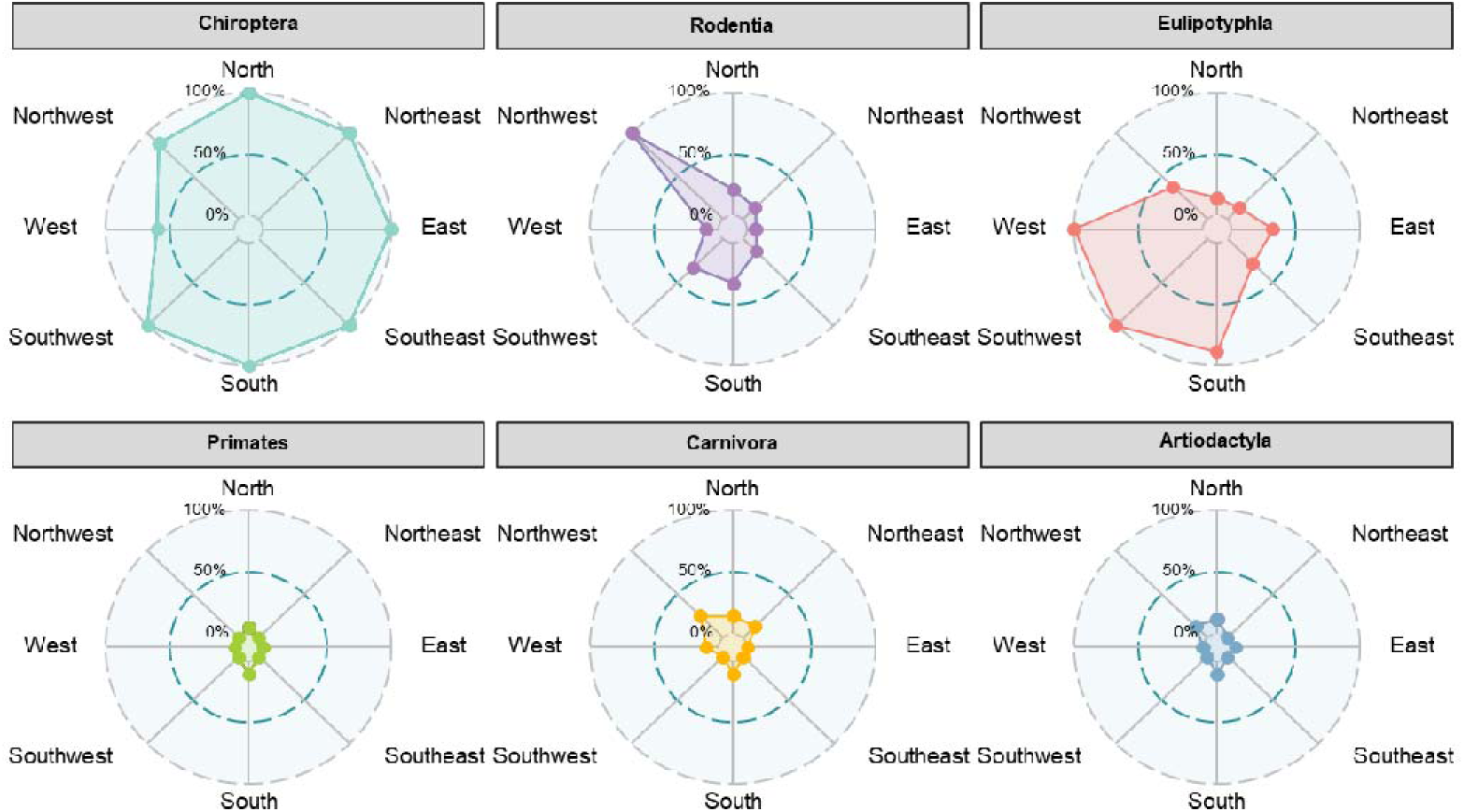
The directions of new mammal records occurred relative to their known distribution ranges across orders (orders with less than four species were not shown).

**Fig. S4.**
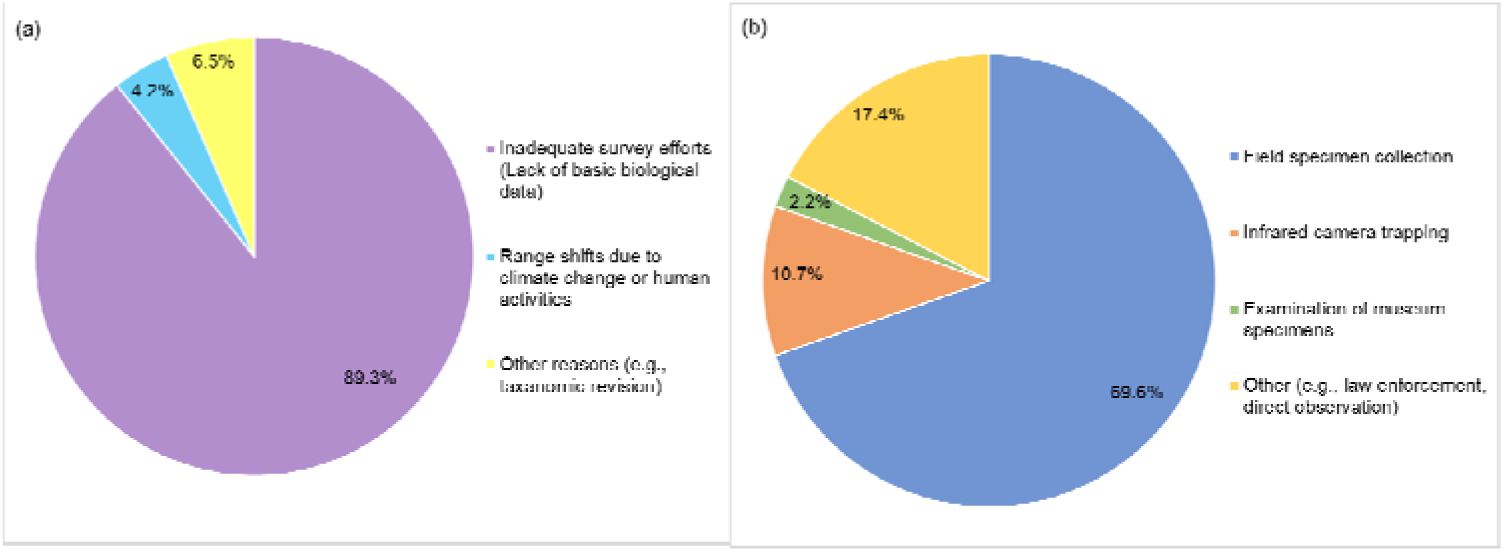
Reasons (a) and methods (b) for discovering new mammal records

