## Supplementary material for "Spatio-temporal distribution patterns and ecological correlates of new mammal records in China": Supporting Information.docx

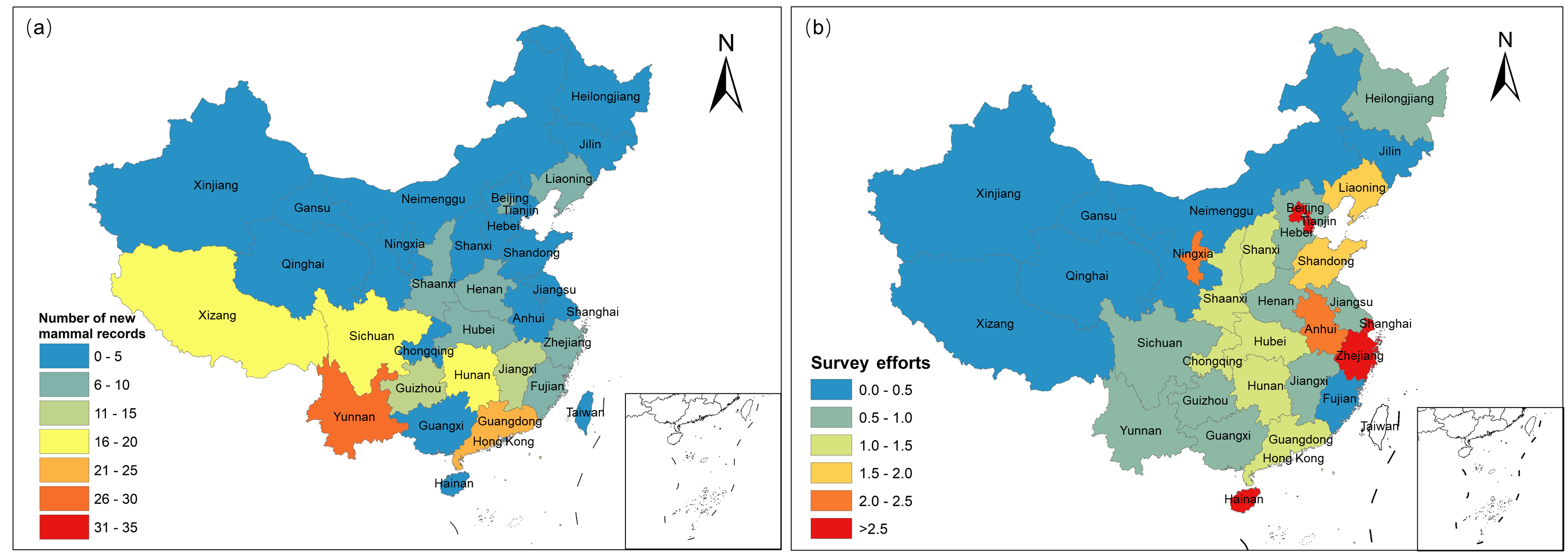

Fig. S1 The spatial patterns of provincial-level new mammal records (a) and survey efforts (b) between 2001 and 2023 in China. Note: the survey effort was calculated as the number of publications targeted for mammal field surveys of each province divided by administrative area and mammal species richness.

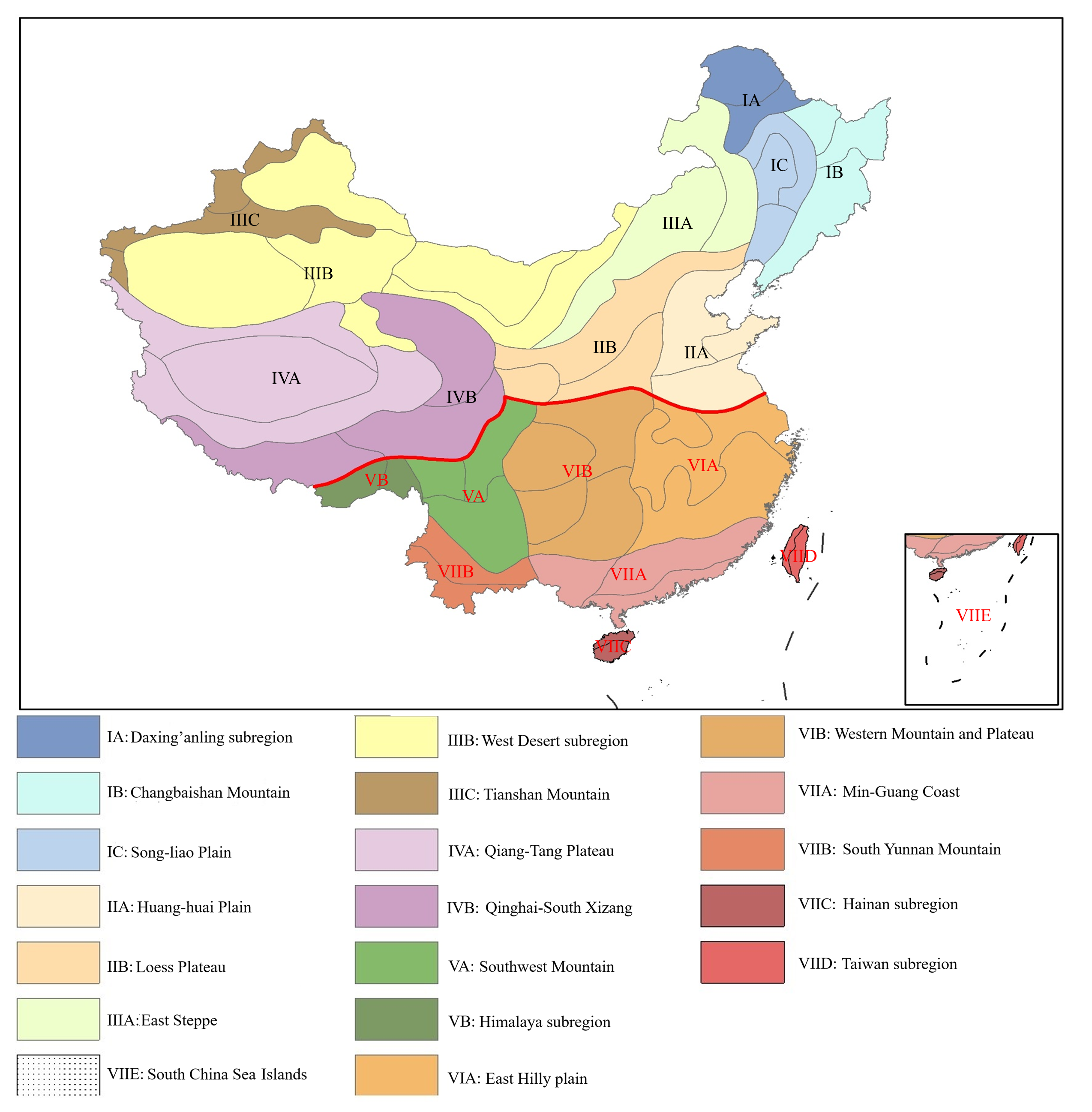

Fig. S2 Map of zoogeographical sub-regions of China for terrestrial vertebrates (Adapted from Gao et al. 2017). The solid red line across it represents the boundary between the Palearctic and Oriental realms.

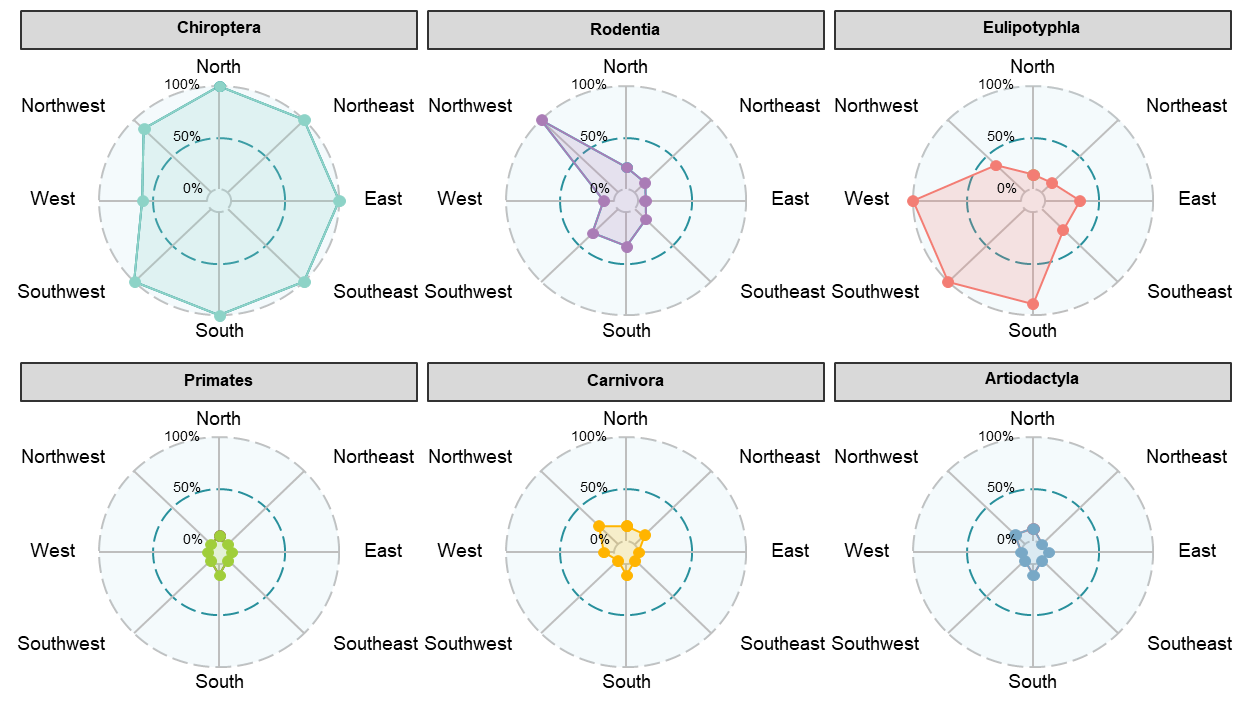

Fig. S3 The directions of new mammal records occurred relative to their known distribution ranges across orders (orders with less than four species were not shown).

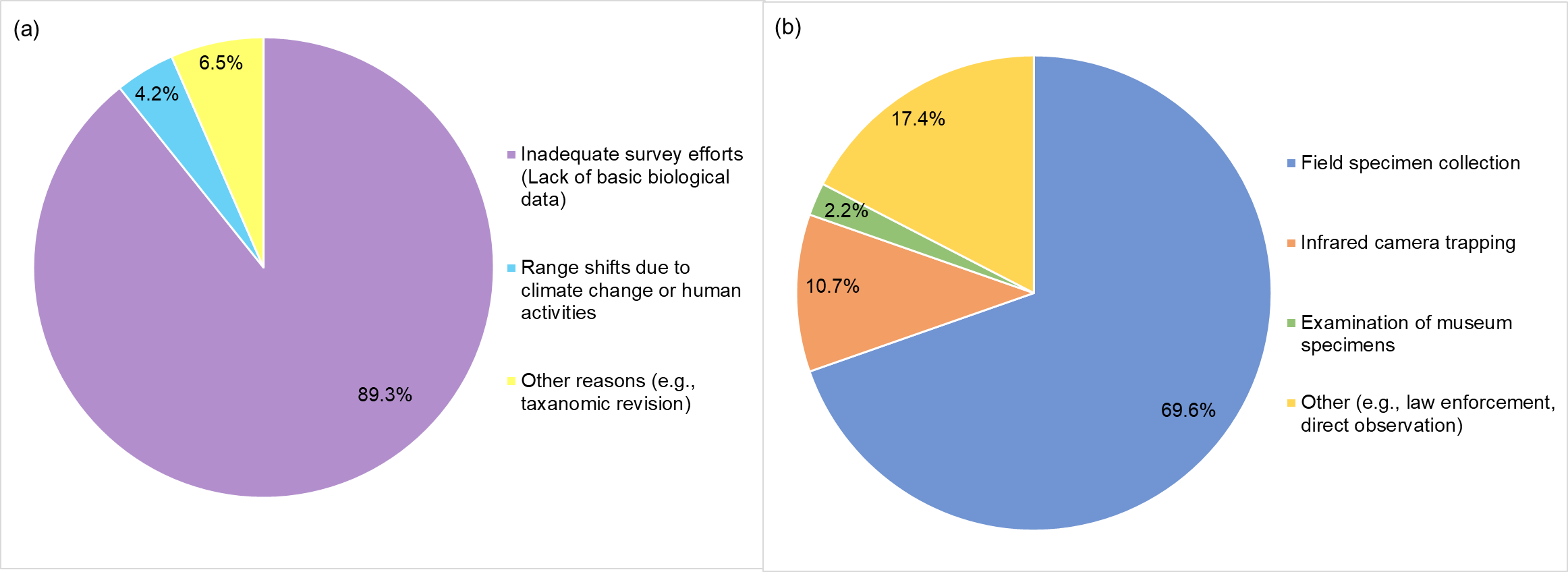

Fig. S4 Reasons (a) and methods (b) for discovering new mammal records
